# Before the brink: sublethal impacts of climate change on stingless bee flight performance

**DOI:** 10.1101/2024.06.17.599256

**Authors:** Carmen Rose Burke da Silva, Lachlan David Macnaughtan, Oliver William Griffith, Ajay Narendra

## Abstract

**Aim:** We aimed to understand the sublethal impacts of climate change on native bee ecology. We evaluated how flight performance, a key trait for predator escape, dispersal, and pollination, is impacted by temperature. We aimed to understand how species geographic ranges shape species thermal performance curves (TPCs) and determine species vulnerability to further warming.

**Location:** Australia

**Time period:** Present

**Major taxa studied:** Stingless bees

**Methods:** We tested the flight speed and acceleration of two species of stingless bees, *Austroplebeia australis* and *Tetragonula carbonaria* at seven test temperatures. We examined how the climate experienced by species throughout their ranges shapes their TPCs. Inferences were made on how sublethal increases in temperature will likely impact key ecological activities associated with flight by estimating the proportion of species geographic ranges that experience temperatures that exceed their thermal optima.

**Results:** Species TPCs reflected the thermal environments they inhabit. *A. australis*, which experiences greater thermal variation and hotter temperatures throughout their range, had a broader TPC and higher thermal optima. However, *A. australis* also had faster flight performance than *T. carbonaria,* rejecting the jack-of-all-trades master-of-none hypothesis. Further increases in temperature will reduce flight performance of *T. carbonaria* at a faster rate (due to their narrower TPCs) than *A. australis*, however, a larger proportion of the *A. australis* range is currently exposed to temperatures above their thermal optima.

**Main conclusions:** Climates will impact species ecology before they reach their lethal limits. Our study supports the idea that species fundamental niche breadths are linked to their geographic ranges, but that trade-offs are not associated with flight performance and ability to perform across a broad range of temperatures. When assessing vulnerability to climate change, it is important to consider how temperature impacts species traits, but also the climates that species experience throughout their unique geographic ranges.

## Introduction

The capacity to tolerate further anthropogenic climate change is often evaluated by using lethal measures of heat tolerance such as upper thermal limits (Kellermann *et al*., 2009; Hoffmann & Sgro, 2018; Bennett *et al*., 2021). These lethal measures can be helpful for ranking species vulnerability to climate change by calculating metrics such as warming or thermal safety margins (da Silva et al., 2023; Sunday et al., 2014). However, climate change will have sublethal impacts on species performance and fitness before they reach their lethal limits (da Silva et al., 2020; van Heerwaarden & Sgrò, 2021), which will likely affect their geographic ranges, and ability to provide functional roles in ecosystems at lower temperatures than predicted from metrics such as CT_MAX_ (maximum critical thermal tolerance) (Jaboor *et al*., 2022; White & Dillon, 2023). Thus, examining sublethal indicators of species responses to climate change is key for understanding how climate change will influence organismal performance, fitness, and ecosystem function in the future.

Understanding how temperature affects ecologically important performance traits, such as flight, can provide insights into species evolutionary histories and how further warming will influence species functional ecology (Clusella-Trullas *et al*., 2011). For most insects, flight is a key trait that facilitates escape from predators, finding mates and resources, and dispersal (Kenna *et al*., 2021). For pollinating insects, such as bees, which are the most important animal pollinators on earth (Michener, 2000), changes in temperature can influence their thermal foraging windows and therefore pollination capabilities (Jaboor *et al*., 2022), which could have a range of negative cascading effects throughout natural and agro-ecosystems.

Estimating the shape of species thermal performance curves (TPC) for traits with clear ecological relevance, such as flight, can provide crucial information on the range of thermal conditions within which species can perform optimally, and how further climate warming will impact species life history traits (Angilletta Jr, 2009; Jayatilaka *et al*., 2011; Kenna *et al*., 2021) (Fig 1). Generally, once a species thermal performance optima (Fig 1) is reached, performance rapidly declines with further increases in temperature (Angilletta Jr, 2009). Understanding how quickly performance declines with temperatures above the thermal optima is thus crucial for understanding sublethal effects of climate change on performance and fitness. The slope of the relationship between performance and temperature post thermal optima depends on whether species have narrow or broad thermal performance breadths (Fig 1), which is hypothesised to depend on species evolutionary histories and geographic ranges (Gilchrist, 1995; Gaston & Spicer, 2001; Calosi *et al*., 2010). For example, species with broad geographic ranges (which tend to experience a great deal of thermal variability throughout their range) are expected to have broad TPCs, and species with narrower geographic ranges (that likely experience less thermal variability throughout their range) are expected to have narrower TPCs (Fig. 1) (Gaston & Spicer, 2001; Calosi *et al*., 2010; Clusella-Trullas *et al*., 2011), however, support for these generalisations is mixed (da Silva et al., 2019).

**Figure 1.**
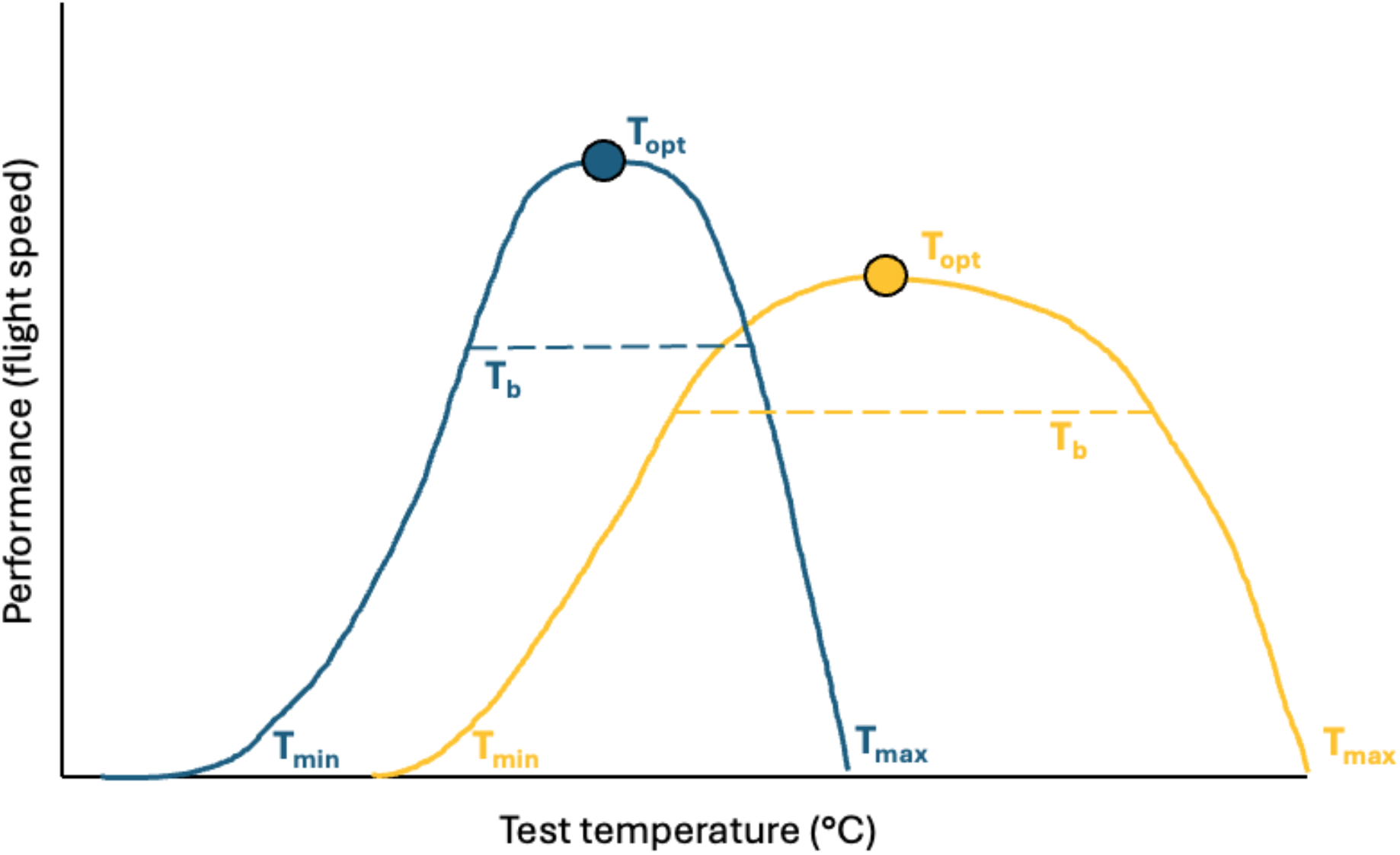
Conceptual illustration of the thermal flight performance curves of a species with a narrow geographic range that experiences a moderate amount of thermal variation (blue), and a species with a very broad geographic distribution that experiences a large degree of thermal variability. T_opt_ = optimal performance temperature, T_b_ = thermal performance breadth, T_min_ = thermal minimum, T_max_ = thermal maximum.

To understand how climate shapes species TPCs and the sublethal impacts of climate change on bee ecology, we examined the flight TPCs of two species of endemic Australian stingless bee, *Austroplebeia australis* and *Tetragonula carbonaria*. *A. australis* has a broad arid/tropical geographic range where they likely experience more climatic variability across their range (Fig 2) (Dollin et al., 2015; Halcroft et al., 2013), than *T. carbonaria* which has a narrower subtropical/temperate distribution (Fig 2). *A. australis* are also likely to experience hotter maximum environmental temperatures (with their tropical/arid range) than *T. carbonaria*, which is reflected by their higher critical thermal maxima (Nacko *et al*., 2023). Stingless bees are generalist foragers that make important contributions to pollination of a wide range of native flora throughout the world’s tropics and subtropics (Bueno *et al*., 2023). They can also be kept in hives and are used as managed pollinators of a variety of fruit and crops in many regions of the world, including Australia (Halcroft, 2013; Heard, 1999).

**Figure 2.**
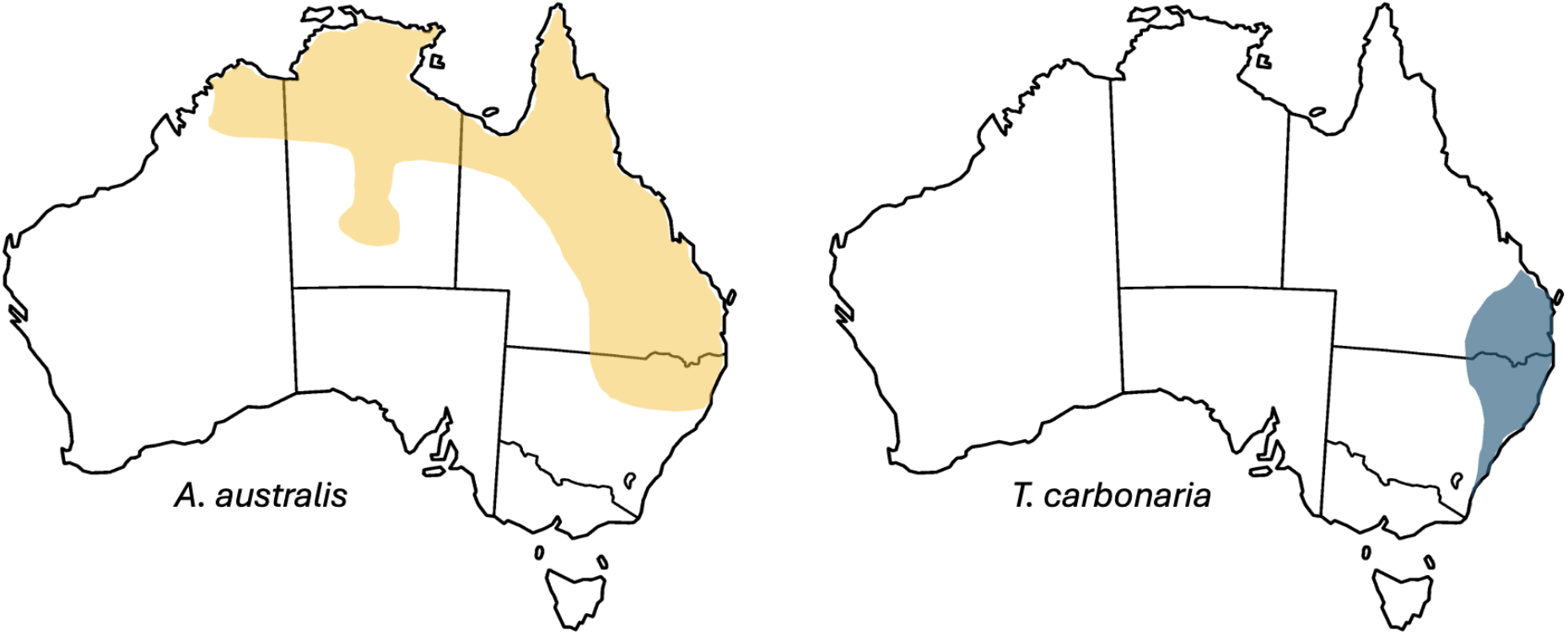
Geographic range of *Austroplebeia australis* (left) based on observed records from (Dollin *et al*., 2015) and *Tetragonula carbonaria* (right) distribution adapted from (Heard, 2016; Vlasich-Brennan, 2023).

We tested the thermal flight performance of *A. australis* and *T. carbonaria* across seven test temperatures. We predicted that the climatic conditions species experience throughout their geographic ranges would explain differences in the shapes of their thermal performance curves (Calosi *et al*., 2010; Clusella-Trullas *et al*., 2011). In particular, we expected *A. australis* would have broader thermal performance curves than *T. carbonaria* due to their broader geographic range. If *T. carbonaria* have a narrower TPC as predicted, their thermal performance would therefore decline more rapidly after their thermal optima is reached with exposure to further warming (i.e. more vulnerable to increases in environmental temperature). We hypothesised that *A. australis,* which has a higher CT_MAX_ (Nacko *et al*., 2023), would have a higher thermal optima than *T. carbonaria* as they experience higher thermal extremes throughout their geographic range. Finally, we predicted *T. carbonaria* to fly faster than *A. australis* at their thermal optima. This prediction was made because species with narrower thermal performance curves are expected to perform better than thermal generalists (jack-of-all-trades master of none hypothesis) due to energetic allocation trade-offs between performance and thermal generalism (Levins, 1968). However, this hypothesis has equivocal support, where the jack-of-all-trades is also known to be a master of all (Huey & Hertz, 1984).

Lastly, to infer how warming climates are already impacting bee thermal ecology, we estimated the proportion of each species geographic distribution where temperatures will exceed species thermal optima, and the upper edge of their thermal breadths (where their performance drops below 80% capacity). To achieve this, we constructed species geographic distribution models based on species occurrence records, pseudo absence points, and climate rasters.

## Methods

### Experimental protocol

*Tetragonula carbonaria* and *Austroplebeia australis* were collected from nest boxes (7 and 3 boxes respectively) on the Macquarie University North Ryde campus, New South Wales Australia. All nests have been located on campus for multiple years, and thus all bees were locally acclimatized to the same environmental conditions. Two of the *A. australis* nests were sourced from https://zabel.com.au/ in 2003 (one black and one orange abdomen phenotype), and the third nest was a black phenotype sourced from southeast Queensland in 2008. The *T. carbonaria* nests were sourced from the Ku-ring-gai council stingless bee program. Bees were collected from nest entrances using a respirator (in batches of 3 individuals per nest multiple times per day) and placed into plastic collection vials with a foam lid to allow airflow. Bees from all nests were placed into an acclimation chamber set at the same temperature as the daily test temperature for one hour prior to testing their flight performance at different acute test temperatures. We tested flight performance at seven acute test temperatures (18, 22, 26, 30, 34, 38, 42 °C) (one test temperature per day) in a randomized order (test day order in Supplementary data). We tested a total of 195 *A. australis* and 210 *T. carbonaria* across all test temperatures where roughly 30 individuals of each species were tested at each test temperature per day (sample sizes for each test temperature provided in Supplementary Table 1). Individuals were tested once and then released at the end of the day. Because worker numbers are so high (∼ 4,000 & 10,000 for *A. australis* and *T. carbonaria* respectively), it is unlikely we re-sampled the same individuals at different test temperatures on different days.

To measure flight performance (speed and acceleration), individual bees were placed at the entrance of a transparent temperature-controlled flight tunnel (1m in length, 10cm in diameter). The flight tunnel was installed within a 46L cooler box which was fitted with a TE Technology AC-027 air cooler/heater (Figure 3). The temperature of the cooler box and flight tunnel was set and maintained with a TE Technology TC-720 temperature controller. Temperature was monitored inside the flight tunnel with a TE Technology MP-3193 thermistor, which communicates with the temperature controller on how to adjust the temperature to maintain the set test temperature. The flight tunnel had many tiny holes drilled into it so that the test temperature would permeate homogeneously throughout the tunnel. The tunnel entrance was fitted with a styrofoam plug to maintain tunnel temperature, which was opened to place bees into the chamber. A thick (6mm) perspex sheet replaced the lid of the cooler box so that flights could be filmed at 100 frames per second using a Chronos 2.1 high speed camera (Kron Technologies, Canada) that was set up on a tripod with a direct dorsal view of the transparent flight tunnel.

**Figure 3.**
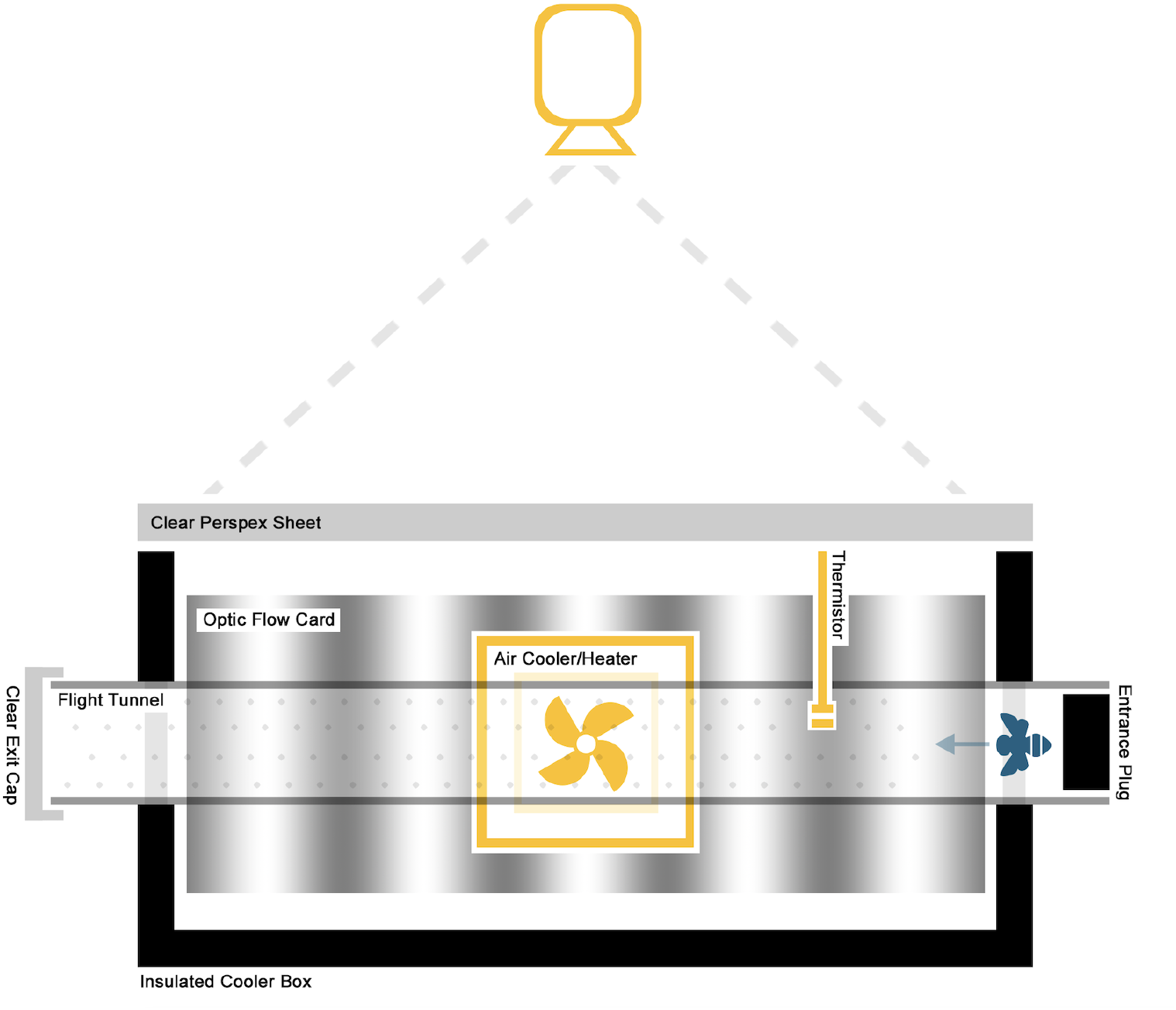
Diagram of temperature-controlled flight tunnel experimental setup.

The end of the flight tunnel was positioned so that bees had the view of the sky as an escape stimulus. A clear cap (top of a large petri dish) was placed at the end of the tunnel so that bees could not escape (and to keep tunnel temperature constant), but could still see the outside escape route. The inner walls of the cooler box were lined with vertical black and white gratings to provide optic flow information. Immediately upon release into the flight chamber bees flew towards the end of the tunnel to attempt escape. Bees were then collected and placed into individual vials. In batches of about 20, bees were then placed in a refrigerator so that they would slow down enough to allow a body mass measurement to be taken on a microbalance (Sartorius Entris II, BCA224I-1S). Bees were released back at their nest boxes once all bees were tested for the day so that the same bee was not tested twice throughout the day, and to avoid killing bees after the experiment.

### Flight performance extraction from video footage

We carried out a frame-by-frame analyses and tracked head position of each bee using the DLTdv8 app (Hedrick, 2008) within Matlab version R2023b (Mathworks, Natick, Massachusetts). We extracted x,y coordinates, converted pixels into centimetres and smoothed flight paths using a smoothing function. Flight speed was extracted by calculating the distance between each consecutive point within the flight path and then using the eq. 1 to calculate speed for each consecutive point comparison.

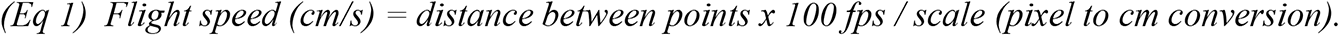

Flight acceleration was calculated using eq. 2.

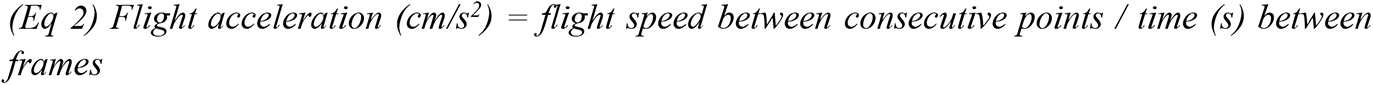

We then extracted the maximum flight speed and acceleration for each individual at each test temperature (flight performance code included within Supplementary Matlab code).

### Statistics

To determine how warming impacts stingless bee flight performance and evaluate how species geographic ranges shape their thermal performance curves, we ran linear mixed effect models using the nlme package (Pinheiro *et al*., 2017) in the statistical program R (R Development Core Team, 2019). We ran two models, one with maximum flight speed and another with maximum acceleration as the response variable. Predictor variables were the same in both models and included body mass, test temperature (as a second-degree polynomial), and species, as well as an interaction between test temperature (as a second-degree polynomial) and species. Colony number was included as a random factor within all models. We used model reduction and comparison using model Akaike Information Criterion (AIC) to arrive at a model that explained the most variation in flight speed and acceleration. Significance of predictor variables were tested using likelihood ratio tests.

### Thermal performance optima and breadth

We calculated the thermal optima and maximum flight performance by estimating the quadratic equation associated with each species thermal performance curve. We then calculated the temperature at which performance was highest (thermal optima) by calculating the x vertex of the quadratic equation using eq. 3,

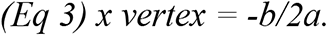

To determine the maximum flight performance at the thermal optima, we calculated the y vertex using eq. 4,

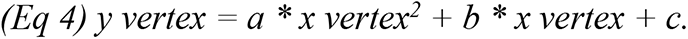

To determine the thermal performance breadths of each species we calculated the range of test temperatures where performance was within 80% of the thermal performance optima.

### Climatic variability throughout species geographic ranges

To quantify the amount of climatic variability within each species geographic range, we extracted their observed GPS coordinates from GBIF (doi: https://doi.org/10.15468/dl.6refur & doi: https://doi.org/10.15468/dl.gmfphe). *T. carbonaria* is known to have isolated allopatric populations in Queensland, apparently restricted to higher altitude ranges of Australia’s Great Dividing Range. It also has a morphologically cryptic cogeneration (*T. hockingsi*) which occurs in many lower altitude regions throughout Queensland, making it challenging to precisely map the distribution in some areas. For our models, we therefore chose a conservation estimate of *T. carbonaria* distribution by restricting only to (i) southern populations and (ii) regions with more than one datapoint. From these coordinates we extracted the maximum (BIO5) and minimum (BIO6) temperature, and mean annual precipitation (BIO12) at each occurrence record for each species from Worldclim2.1 at 2.5 minute resolution (Fick & Hijmans, 2017).

### Predict impact of climate warming on each species

To determine the overall rate at which performance will decline in each species as environmental temperatures surpass their thermal optima, we estimated the linear slopes between each species’ thermal optima and their performance at 42 °C (hottest test temperature).

To understand how climate change is already impacting flight performance, we calculated the proportion of each species range where hottest temperatures of the hottest month (BIO5) is higher than the thermal optima and upper thermal breadth for each species. For this, we constructed generalised linear species distribution models following the online tutorial by Jeff Oliver (https://jcoliver.github.io/learn-r/011-species-distribution-models.html), which included creating pseudo absence data, training our distribution models, and predicting species ranges based off a likelihood threshold.

## Results

### Flight speed and acceleration

For both the flight speed and acceleration models, we found that TPCs differed between the two species, where the best fitting models included a significant interaction between test temperature and species (flight speed: F = 14.0, df = 2/390, *p* < 0.001; acceleration: F = 15.62, df = 2/390, *p* < 0.001) (Table 1 & 2). *A. australis* had a higher optimal performance temperature, a broader thermal performance curve, and greater optimal flight performance for both maximum speed and acceleration compared to *T. carbonaria* (Table 3; Figure 4). Flight performance (speed and acceleration) modelled 95% confidence intervals around the mean overlapped at temperatures below the 30 °C suggesting no difference in flight performance at colder test temperatures, however, from 30 °C onwards, *A. australis* flew faster and accelerated faster than *T. carbonaria* on average.

**Figure 4.**
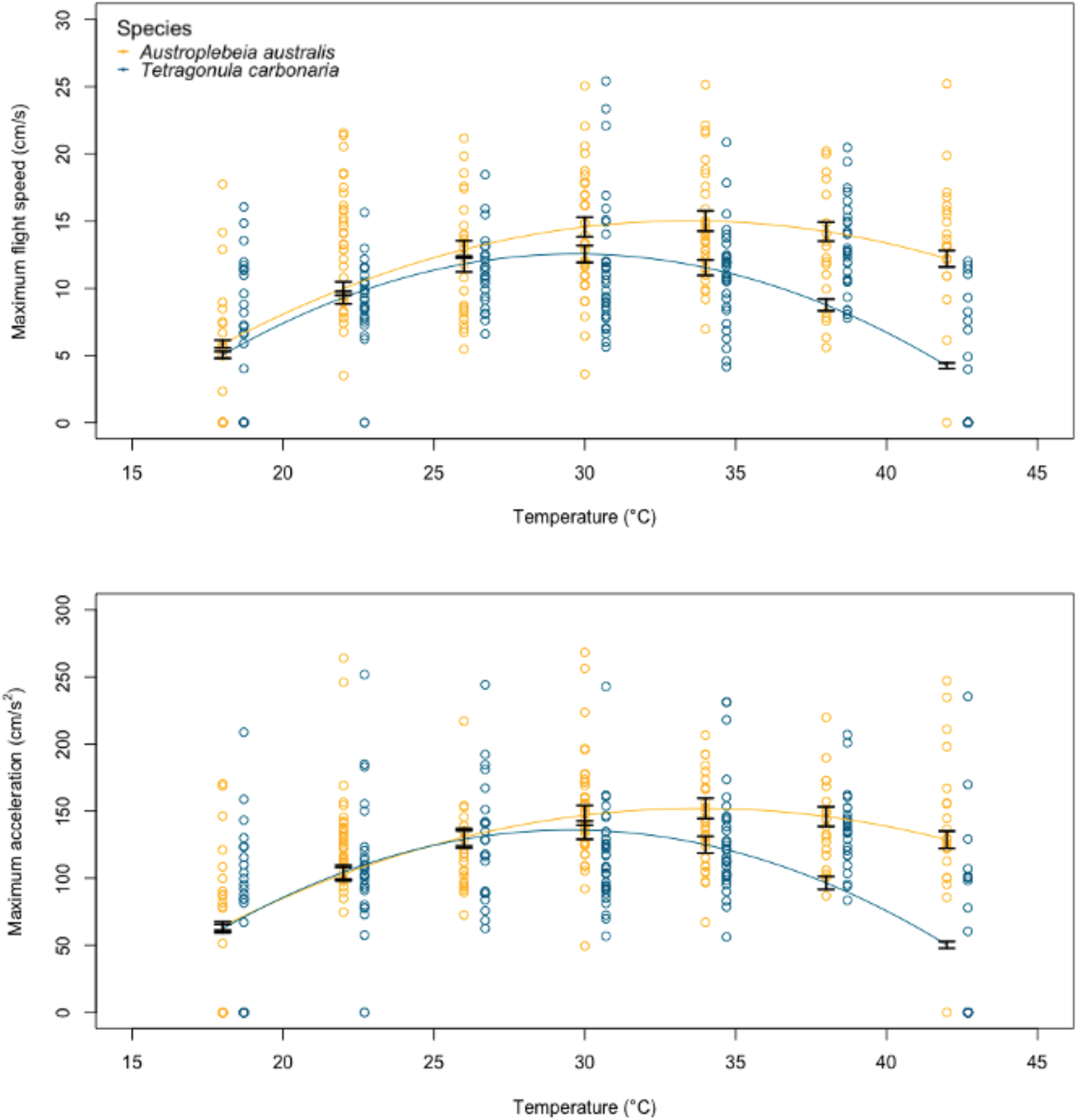
Maximum flight speed (top) and acceleration (bottom) of *A. australis* (yellow) and *T. carbonaria* (blue) across the 7 experimental test temperatures. Lines go through the model means of each species performance at each test temperature. 95% confidence intervals are represented by error bars around the modelled mean performance for each species at each test temperature.

**Table 1.** AIC model comparison for maximum flight speed and maximum acceleration.

| Model | df | AIC | ΔAIC |
| --- | --- | --- | --- |
| *Maximum flight speed* |  |  |  |
| poly(Test temp,2) * species | 8 | 2406.19 | 0 |
| poly(Test temp,2) + species | 6 | 2429.39 | 23.2 |
| Mass + poly(Test temp, 2) * species | 44 | 2446.76 | 40.57 |
| Mass + poly(Test temp, 2) + species | 42 | 2467.68 | 61.49 |
| Mass + Test temp * species | 42 | 2558.12 | 151.93 |
| *Maximum acceleration* |  |  |  |
| poly(Test temp,2) * species | 8 | 4292.02 | 0 |
| poly(Test temp,2) + species | 6 | 4318.54 | 26.52 |
| Mass + poly(Test temp, 2) * species | 44 | 4334.71 | 42.69 |
| Mass + poly(Test temp, 2) + species | 42 | 4355.94 | 63.92 |
| Mass + Test temp * species | 42 | 4432.61 | 140.59 |

**Table 2.**
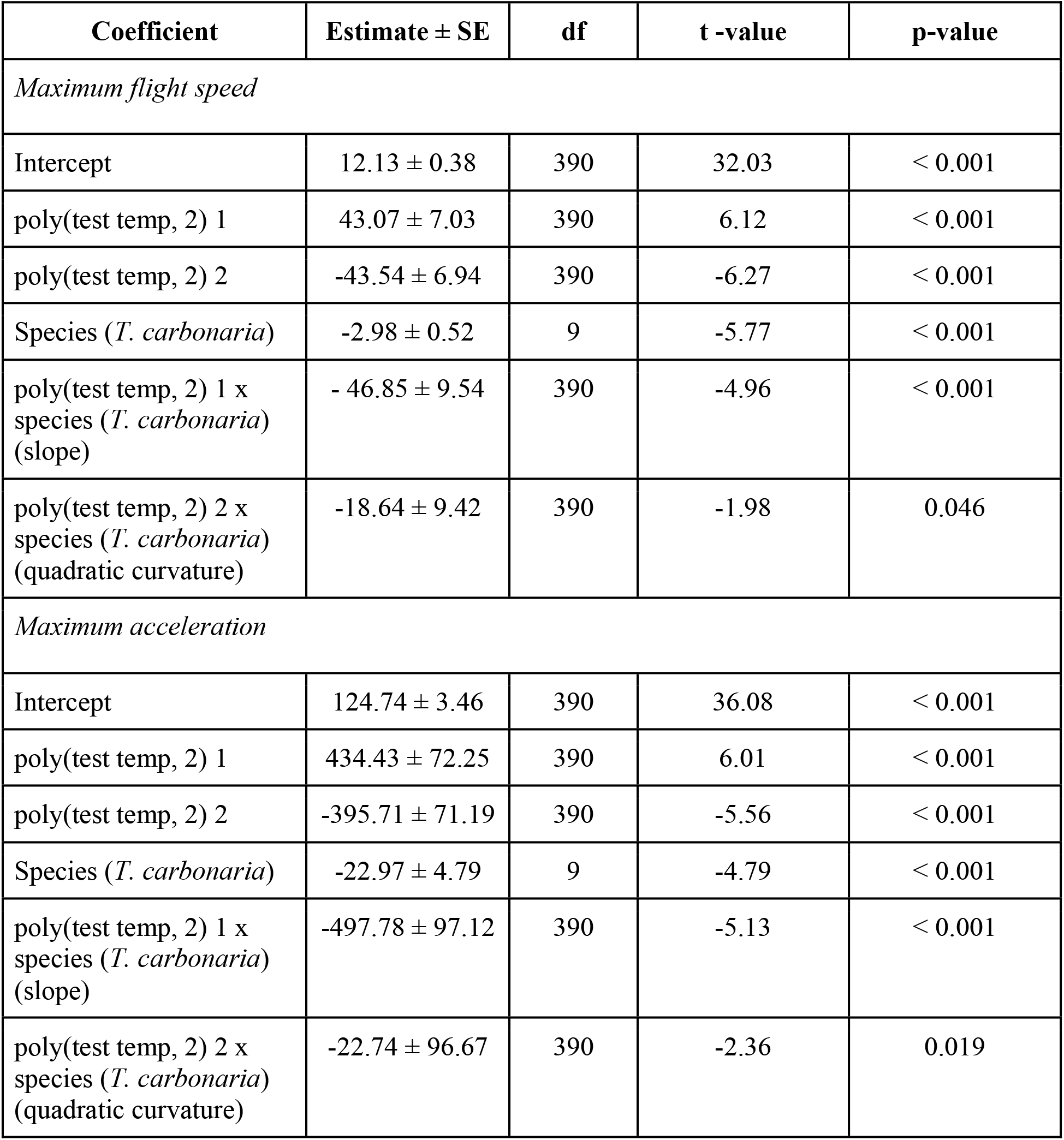
Model summary of the effect of test temperature (modelled as a second-degree polynomial) on maximum flight speed for each species.

**Table 3.**
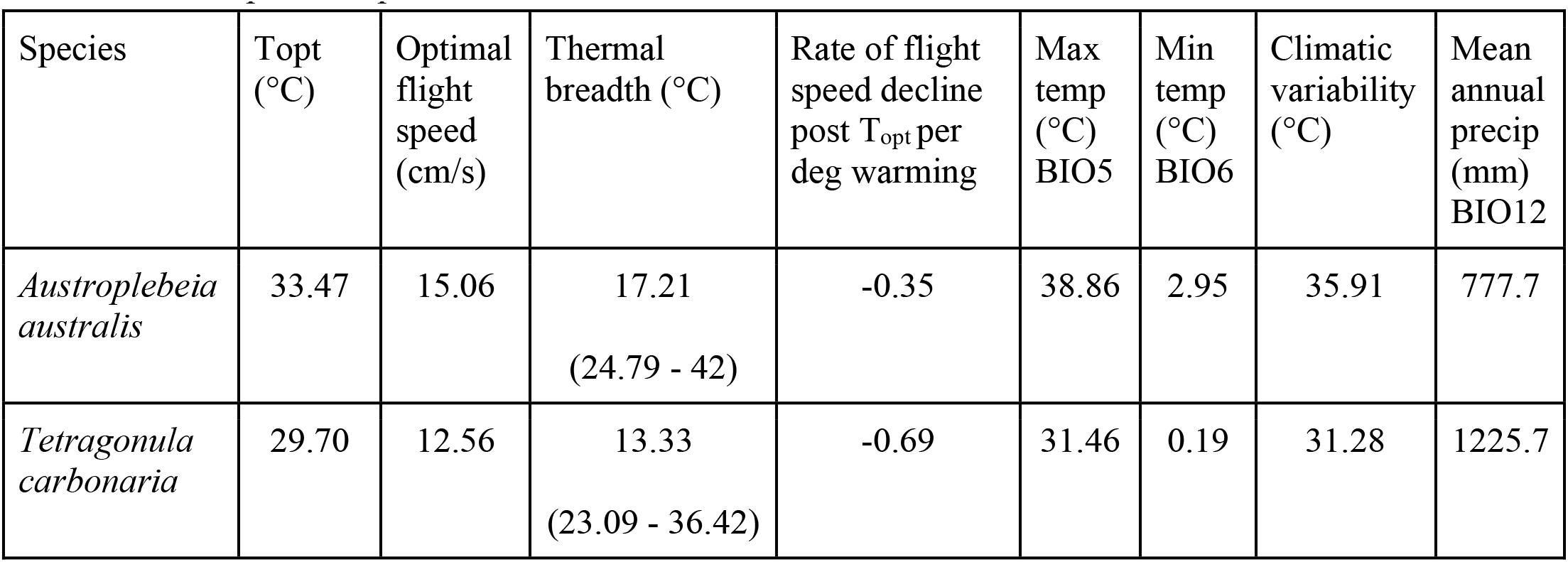
Summary table of species thermal flight speed performance curve metrics and climatic conditions each species experiences at their occurrence records.

Body mass was not included within the model with the lowest AIC and it did not have a significant effect on flight speed or acceleration so it was removed as a predictor variable so that flight speed of all individuals (including those that escaped after the flight speed testing and could not be weighed) could be included within the analysis.

To understand the impact of warming on each species, we estimated the rate at which flight speed performance declines once each species thermal optima is reached. We found that flight speed performance declines almost twice as rapidly with each degree increase in environmental temperature in *T. carbonaria* than *A. australis* (Table 3). We also quantified the climatic conditions (environmental temperature (minimum, maximum, thermal variability) and mean precipitation) each species experiences at their occurrence records. Surprisingly, the amount of thermal variability that *A. australis* and *T. carbonaria* experiences throughout their range is not as different as we would have expected based on differences in their geographic range extents. However, *A. australis* does experience greater thermal variability, higher maximum environmental temperatures, and they live in drier habitats than *T. carbonaria* overall (Table 3).

To evaluate how warming climates are impacting species performance, we estimated the proportion of their ranges where environmental temperatures already exceed their thermal optima and the upper edge of their thermal breadths (Table 4). We found that BIO5 temperatures (hottest temperatures of the hottest month) will not exceed the upper edge of species thermal breadths, meaning that flight performance will not drop below 80% of their maximum flight capacity throughout their entire range (Figure 5). However, very large proportions of species ranges are already experiencing environmental temperatures (BIO5) above their thermal optima (74.15, and 59.39% for *A. australis* and *T. carbonaria* respectively), where flight performance will decline with further increases in environmental temperature, and the rate of decline will be more rapid in *T. carbonaria* than *A. australis* (Tables 3 & 4).

**Figure 5.**
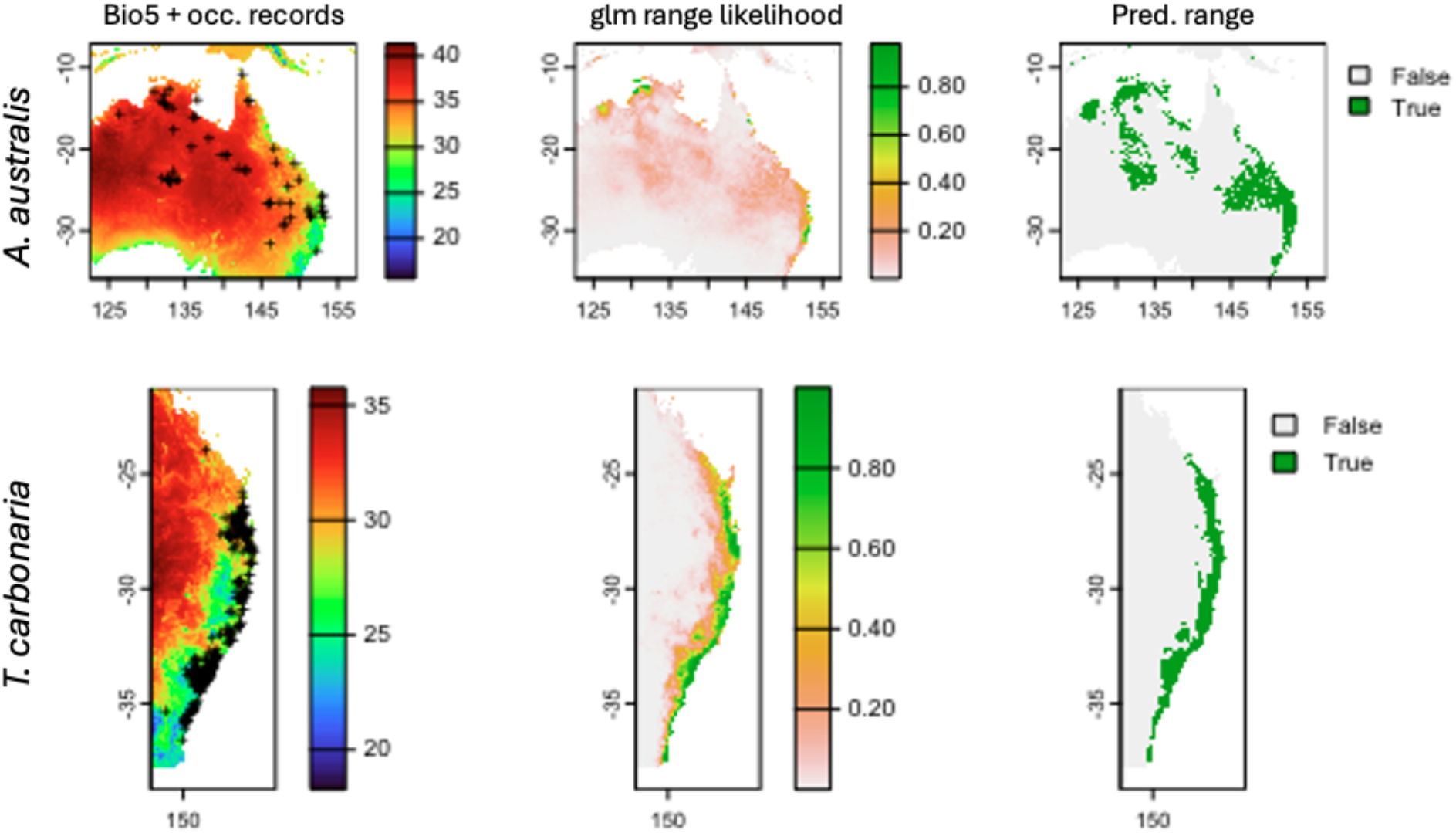
Geographic distribution of *A. australis* and *T. carbonaria*. Left panel: Species occurrence records plotted over Worldclim variable BIO5 climate raster (hottest temperature of the hottest month) for *A. australis* (top row) and *T. carbonaria* (bottom row). Middle panel: generalised linear species distribution model predictions of species geographic range where colours indicate likelihood of occurrence. Right: predicted species geographic ranges above modelled likelihood threshold values (true (green) indicates where species are highly likely to occur, false (grey) areas indicate where species are unlikely to occur). BIO5 values within species predicted ranges were used to estimate the proportion of species ranges that exceed species T_opt_ and upper edge of their thermal breadth.

**Table 4.** Percentage of species’ distribution ranges where BIO5 (hottest temperature of hottest month) is greater than each species upper thermal breadth (where performance dips below 80% capacity), or is over T_opt_ under present climatic conditions.

| Species | BIO5 > $T_{opt}$ | BIO5 > upper Tb |
| --- | --- | --- |
| *Austroplebeia australis* | 74.15% | 0% |
| *Tetragonula carbonaria* | 59.39% | 0% |

## Discussion

Using a custom-made flight tunnel, we investigated how escape flight performance (speed and acceleration) varied with ecologically relevant test temperatures in two species of stingless bee, *A. australis* and *T. carbonaria*. We found that both species have broad thermal breadths where they can maintain flight speed within 80% of their capacity across thermal ranges of 13 °C (23.09 - 36.42 °C) and 17 °C (24.79 - 42 °C), respectively. These thermal flight performance breadths are quite broad compared to other insect species, such as 11 species of broadly distributed *Drosophila* which only have thermal flight performance breadths between ∼ 3 - 7 °C (De Araujo *et al*., 2019). The broad thermal flight performance breadths of Australian stingless bees are likely to facilitate pollination across broad climatic conditions, and potentially buffer pollination services for the plants they pollinate in the near-term as climates continue to change.

We examined how geographic range influences species’ thermal flight performance curves. We found that *A. australis* which has a broad arid/tropical geographic range had a broader thermal performance breadth for both flight speed and acceleration than *T. carbonaria,* which has a narrower subtropical/temperate range. This finding supports the hypothesis that species that experience more climatic variation throughout their geographic ranges should have broader thermal tolerances and breadths (Gilchrist, 1995; Gaston & Spicer, 2001; Calosi *et al*., 2010; Clusella-Trullas *et al*., 2011). This hypothesis is supported by both meta-analyses and experimental research (Calosi *et al*., 2010; Rohr *et al*., 2018). However, there are many exceptions to the rule, where sometimes species that experience large degrees of thermal variability evolve narrow thermal performance breadths, but have a great capacity to shift their thermal performance via seasonal acclimation, particularly when they live in environments with predictable seasonal variation (da Silva et al., 2019).

We also found that *A. australis* had a higher thermal performance optima (33.47 °C) than *T. carbonaria* (29.70 °C), reflecting the higher maximum environmental temperatures that they experience throughout their geographic range. However, speed and acceleration at cooler test temperatures (18 °C - 26 °C) did not differ between the species (overlapping confidence intervals), even though *T. carbonaria* experiences cooler climates on average, likely reflecting the similar absolute minimum thermal environments that they experience throughout their range (Table 3).

The ‘jack-of-all-trades, master-of-none’ hypothesis predicts that species that have broader thermal performance curves should have reduced performance at their thermal optima than thermal specialists, according to the Principle of Allocation, which states that there should be trade-offs between performance and thermal generalism (Levins, 1968). Despite their broader thermal performance breadth, we found that *A. australis* had greater flight performance (speed and acceleration) at their thermal performance optima, and at all test temperatures higher than 26 °C. This means that *A. australis* is not a ‘jack-of-all-trades, master-of-none’ but is in fact a ‘jack-of-all-trades, master-of-all’, supporting the findings originally tested by Huey & Hertz (1984), and reiterated by Gaston & Spicer (2001). This means that in stingless bees there is unlikely to be a trade-off between performance and thermal generalism. However, while the ‘jack-of-all-trades, master-of-all’ hypothesis is supported (Huey & Hertz, 1984), both species had relatively broad thermal performance curves, and it is not clear why *A. australis* had greater flight performance than *T. carbonaria* at test temperatures over 26 °C. Perhaps *A. australis* have evolved faster flight speeds than *T. carbonaria* due to stronger selection on traits the facilitate worker survival. *A. australis* live in smaller colonies than *T. carbonaria* where they are estimated to have a worker population of only 4,000 individuals (Halcroft et al., 2011), compared to 10,000 for *T. carbonaria* (Heard, 2016). There could be stronger selection on traits, such as escape speed and acceleration, that facilitate survival from predators in species that live in smaller groups. When the number of workers bringing resources back to the nest are reduced, loss of workers could have a larger impact on colony fitness. Concordantly, *A. australis* have been reported to have the longest longevity compared to all other eusocial bees recorded (Halcroft *et al*., 2013a), further indicating they have likely developed traits to facilitate survival over long durations compared to other species.

Body mass did not explain differences in flight performance between the two stingless bee species, even though mass was an important predictor for flight speed in different beetle species (Farisenkov *et al*., 2020). However, bumble bees, which are much larger than stingless bees (by about 10x), flew much faster than the stingless bees in our study (Kenna et al., 2021). Perhaps the relationship between body mass and flight speed only becomes apparent with larger differences in body mass. In contrast to our study, Kenna et al., (2021) found no relationship between flight speed and test temperature in bumble bees. They measured flight speed using a flight mill while also assessing how temperature impacts the duration and distance bumble bees flew (where they did find that endurance and distance have a ‘performance curve shape’ relationship with temperature). The absence of a relationship between flight speed and temperature in their study was likely because they did not measure escape speed per se, but mean and maximum speed over long distances where bees will need to pace themselves. Our study demonstrates that escape flight speed and acceleration has a very clear relationship with temperature in Australian stingless bees.

### Impacts of climate change on stingless bees

To quantify the vulnerability of these important pollinators to climate change, we estimated the rate at which flight performance will decline when environmental temperature surpasses their thermal optima. We found that flight speed and acceleration will drop at a faster rate in *T. carbonaria* than *A. australis*, owing to their narrower thermal performance curves. However, when assessing vulnerability to climate change it is also important to consider differences in the environmental conditions species experience within their ranges. Because *T. carbonaria* inhabit climatic conditions that are cooler than *A. australis* on average, neither species is exposed to (mean) maximum temperatures higher than the upper edge of their thermal performance breadth under current climate conditions, which means they can maintain flight performance within 80% capacity throughout their entire geographic ranges. However, we also estimated the proportion of each species range that experiences environmental temperatures over each species thermal optima. These calculations show that a large proportion of each species range is already exposed to environmental conditions above their thermal optimum. However, even though *T. carbonaria* have a lower thermal optima to *A. australis*, a greater proportion of the geographic range of *A. australis* will be exposed to environmental temperatures above their thermal optima because they inhabit environments that are much hotter than *T. carbonaria* on average (Table 4; Figure 5).

Escape flights need to be fast and precise since the penalty for slow escape is usually severe. Reduced escape speed and acceleration at environmental temperatures above species thermal optima could make stingless bees more susceptible to predation as climates continue to warm. Temperatures above species thermal optima could also impact species agility during escape responses, potentially further impacting their capacity to escape predation and navigate through complex habitats (Card, 2012; Singh *et al*., 2024). However, increased environmental temperatures can also affect locomotory strategies of predators, and it is important to consider how climate change will impact predator-prey relationships (Grigaltchik *et al*., 2012). A study that examined thermal locomotor performance in predator and prey freshwater fish found that at warm test temperatures escape speed of prey species increased and the number of predator attacks decreased, suggesting that predation pressure will decrease at higher temperatures (Grigaltchik *et al*., 2012). Similarly, a recent meta-analysis (comparing species CT_MAX_) found that top predators tend to be more vulnerable to warming climates than primary consumers and pollinators (da Silva *et al*., 2023). Thus, even though stingless bee flight escape responses will be reduced with warming climates, climate change might have an even larger impact on species that predate stingless bees, such as birds, spiders, ants, and centipedes (Rao *et al*., 2008; Vijayakumar *et al*., 2012; Ostwald *et al*., 2018; Roubik, 2023). However, carefully designed experiments are required to examine how warmed environmental temperature impacts predation success by these predatory species.

### Study limitations

In this study, we make inferences on how the climates species experience throughout their geographic ranges shapes their geographic ranges. While we find strong evidence suggesting that a species with a larger geographic range has a broader thermal performance curve, higher thermal optima, and greater performance at warm test temperatures, it is difficult to make generalisations on how climate shapes species thermal performance curves from a study with a species sample size of two. In the future we aim to test the thermal flight performance curves of many species of Australian bee that occur over broad climatic gradients.

We tested the hypothesis that species with broader geographic ranges (which experience greater climatic variation) should have broader TPCs than species with narrow geographic ranges. However, it is worth acknowledging that populations within species might have differently shaped TPCs due to local adaptation (Sinclair *et al*., 2016). While there are very few examples of studies that compare population TPCs across climatic gradients, there are many studies that show thermal limits are correlated with climatic gradients across populations suggesting local adaptation (Castañeda *et al*., 2015; Pereira *et al*., 2017; Healy *et al*., 2019). Because we do not know the original source locations for the *A. australis* in our study we are unable to make inferences on how source local climate might have shaped their TPCs.

To assess the vulnerability of Australian stingless bees to current environmental conditions we quantified the proportion of species ranges where climates exceed their thermal optima and upper edge of their thermal performance breadths. However, these calculations do not account for changes in behaviour which could influence species activity times and the environmental temperatures that species are exposed to. For example, on hot days, stingless bees might not leave their nests, and therefore may not be exposed to extreme thermal environments where their flight performance will be reduced. However, changes in species behaviour, such as activity times also have the potential to negatively impact pollination services throughout ecosystems (Jaboor *et al*., 2022).

### Conclusion

Our data highlight that species thermal performance curves are shaped by the climatic conditions that they inhabit (Calosi *et al*., 2010; Clusella-Trullas *et al*., 2011), and reject the Principle of Allocation hypothesis which states that there are trade-offs between performance and thermal generalism (Levins, 1968; Huey & Hertz, 1984). Our models predict that flight performance will decline with warming climates at a more rapid rate in *T. carbonaria* than *A. australis* due to their narrower thermal performance curves. However, a greater proportion of the range of *A. australis* experiences extreme climates that exceed their thermal optima than *T. carbonaria*.

Global databases of species upper limits, such as GlobTherm (Bennett *et al*., 2018) are useful to understand how climate shapes thermal tolerances (Bennett *et al*., 2021) and rank species vulnerabilities to climate change (Sunday *et al.,* 2014; da Silva *et al.,* 2023). However, to predict the sublethal impacts of climate change on species functional traits, and how these traits might impact the functioning of ecosystems with only moderate climate warming (e.g. impacts of climate change on pollination), similar datasets for species (and populations) thermal performance curves should be collated. Our data shows that stingless bee flight performance will be impacted before environments are so warm that they are unable to function, which could influence their ability to escape from predators, provide pollination services, disperse, and find mates.

## Data Availability Statement

Data will be made freely available upon acceptance on a freely accessible server such as Dryad or equivalent.

## Author contributions

CRBdS conceptualised the project, acquired the funding, did the statistical analyses, and wrote the first draft of the manuscript. CRBdS and LM conducted the experiment. LM extracted the flight paths from video files and contributed to manuscript editing. OG contributed towards writing and editing. AN provided access to a high-speed camera, provided expertise on flight path extraction and analysis, and contributed to writing and editing.

## Acknowledgements

We thank Rosalyn Gloag for providing comments on an early draft of the manuscript.

---

**Supplementary Table 1.** Summary table of *Austroplebeia australis* and *Tetragonula carbonaria* maximum flight speed sample sizes at each test temperature, and the predicted mean flight speeds at each test temperature.

| Test temperature (°C) | n | Mean pred. max flight speed (cm/s) | Mean pred. max. acceleration (cm/s<sup>2</sup>) |
| --- | --- | --- | --- |
| *A. australis* |  |  |  |
| 18 | 29 | 5.83 | 64.34 |
| 22 | 33 | 9.98 | 103.06 |
| 26 | 26 | 12.89 | 130.58 |
| 30 | 36 | 14.56 | 146.91 |
| 34 | 32 | 15.01 | 152.03 |
| 38 | 19 | 14.22 | 145.95 |
| 42 | 20 | 12.2 | 128.67 |
| *T. carbonaria* |  |  |  |
| 18 | 29 | 5.03 | 62.41 |
| 22 | 27 | 9.3 | 104.52 |
| 26 | 24 | 11.8 | 128.99 |
| 30 | 35 | 12.55 | 135.81 |
| 34 | 31 | 11.53 | 124.98 |
| 38 | 28 | 8.76 | 96.51 |
| 42 | 36 | 4.22 | 50.39 |

